# Potent in vitro Neutralization of SARS-CoV-2 by Hetero-bivalent Alpaca Nanobodies Targeting the Spike Receptor-Binding Domain

**DOI:** 10.1101/2021.02.02.429311

**Authors:** Huan Ma, Weihong Zeng, Xiangzhi Meng, Xiaoxue Huang, Yunru Yang, Dan Zhao, Peigen Zhou, Xiaofang Wang, Changcheng Zhao, Yong Sun, Peihui Wang, Huichao Ou, Xiaowen Hu, Yan Xiang, Tengchuan Jin

**Author notes:** To whom correspondence should be addressed: Prof. Tengchuan Jin: Division of Life Sciences and Medicine, University of Science and Technology of China, Hefei, 230027, China; Prof. Yan Xiang: Department of Microbiology, Immunology and Molecular Genetics, University of Texas Health Science Center at San Antonio, San Antonio, Texas, USA. These authors contributed equally to this work (Huan Ma, Weihong Zeng).

## Abstract

Cell entry by SARS-CoV-2 requires the binding between the receptor-binding domain (RBD) of the viral Spike protein and the cellular angiotensin-converting enzyme 2 (ACE2). As such, RBD has become the major target for vaccine development, while RBD-specific antibodies are pursued as therapeutics. Here, we report the development and characterization of SARS-CoV-2 RBD-specific V_H_H/nanobody (Nb) from immunized alpacas. Seven RBD-specific Nbs with high stability were identified using phage display. They bind to SARS-CoV-2 RBD with affinity K_D_ ranging from 2.6 to 113 nM, and six of them can block RBD-ACE2 interaction. The fusion of the Nbs with IgG1 Fc resulted in homodimers with greatly improved RBD-binding affinities (K_D_ ranging from 72.7 pM to 4.5 nM) and nanomolar RBD-ACE2 blocking abilities. Furthermore, fusion of two Nbs with non-overlapping epitopes resulted in hetero-bivalent Nbs, namely aRBD-2-5 and aRBD-2-7, with significantly higher RBD binding affinities (K_D_ of 59.2 pM and 0.25 nM) and greatly enhanced SARS-CoV-2 neutralizing potency. The 50% neutralization dose (ND_50_) of aRBD-2-5 and aRBD-2-7 was 1.22 ng/mL (∼0.043 nM) and 3.18 ng/mL (∼0.111 nM), respectively. These high-affinity SARS-CoV-2 blocking Nbs could be further developed into therapeutics as well as diagnosis reagents for COVID-19.

**Importance:** To date, SARS-CoV-2 has caused tremendous loss of human life and economic output worldwide. Although a few COVID-19 vaccines have been approved in several countries, the development of effective therapeutics including SARS-CoV-2 targeting antibodies remains critical. Due to their small size (13-15 kDa), highly solubility and stability, Nbs are particularly well suited for pulmonary delivery and more amenable to engineer into multi-valent formats, compared to the conventional antibody. Here, we report a serial of new anti-SARS-CoV-2 Nbs isolated from immunized alpaca and two engineered hetero-bivalent Nbs. These potent neutralizing Nbs showed promise as potential therapeutics against COVID-19.

## Introduction

Coronavirus disease 2019 (COVID-19) caused by SARS-CoV-2 has resulted in tremendous health and economic losses worldwide. SARS-CoV-2 belongs to the betacoronavirus genus, which include two other significant human pathogens, the Severe Acute Respiratory Syndrome (SARS-CoV-1) virus and the Middle East Respiratory Syndrome (MERS) virus, first emerging in humans in 2002 and 2012, respectively [1-4]. Currently, several COVID-19 vaccines have been approved for emergency usages by several countries [5, 6]. Remdesivir [7] and dexamethasone [8] have also been approved for treating COVID-19 under emergency use authorization. To more effectively combat COVID-19 and prepare for possible future pandemics, it remains essential to develop new effective drugs targeting coronaviruses.

Virus-specific antibody responses can be readily detected in sera of COVID-19 patients [9-12], and a series of monoclonal antibodies (mAbs) that neutralize SARS-CoV-2 have been isolated from infected individuals [13-18]. Both convalescent plasma and mAbs targeting SARS-CoV-2 have shown promise as therapeutics for treating COVID-19 patients [19-21]. In addition to the conventional mAbs, a distinct type of antibody fragment derived from camelid immunoglobulins, termed V_H_H or nanobody (Nb), is an attractive alternative for COVID-19 treatment. Compared to the conventional antibody, V_H_H is cheaper to produce, has an enhanced tissue penetration, and is more amenable to engineering into multivalent and multi-specific antigen-binding formats [22]. Moreover, Nbs are particularly well suited for pulmonary delivery because of their small size (13-15 kDa), highly solubility and stability [23, 24].

Cell entry by SARS-CoV-2 requires the interaction between the RBD of the viral Spike protein and the receptor ACE2, which is also the receptor for SARS-CoV-1[25-29]. The RBD of SARS-CoV-2 binds to ACE2 with a K_D ,_of ∼15 nM, which about 10-to 20-fold better than that for SARS-CoV-1 RBD[30].

In this study, we report the development and characterization of seven anti-RBD Nbs isolated from alpcacas immunized with SARS-CoV-2 RBD. Furthermore, two high-affinity hetero-bivalent Nbs were developed by fusing two Nbs with distinct epitopes, resulting in antibodies with strong SARS-CoV-2 neutralizing potency.

## Results

### Isolation of anti-SARS-CoV-2 RBD nanobodies from immunized alpacas

Our aim was to develop potent SARS-CoV-2 neutralizing antibodies with favorable biological characteristics. Towards this goal, we immunized two alpacas 3 times with highly purified recombinant SARS-CoV-2 RBD (**Fig. S1**). Total RNA was extracted from 1 × 10^7^ PBMCs from the immunized alpacas and used as the template for synthesizing cDNA. The V_H_H coding regions were amplified from the cDNA and cloned into a phagemid vector, generating a library with about 1.6 × 10^7^ independent clones. Phages displaying V_H_H were prepared from the library with the helper phage and selected with SARS-CoV-2 RBD via two rounds of biopanning. Titration of the output phages after each round of panning indicated that the RBD-binding phages were effectively enriched (**Fig. 1A)**.

**Fig. 1.**
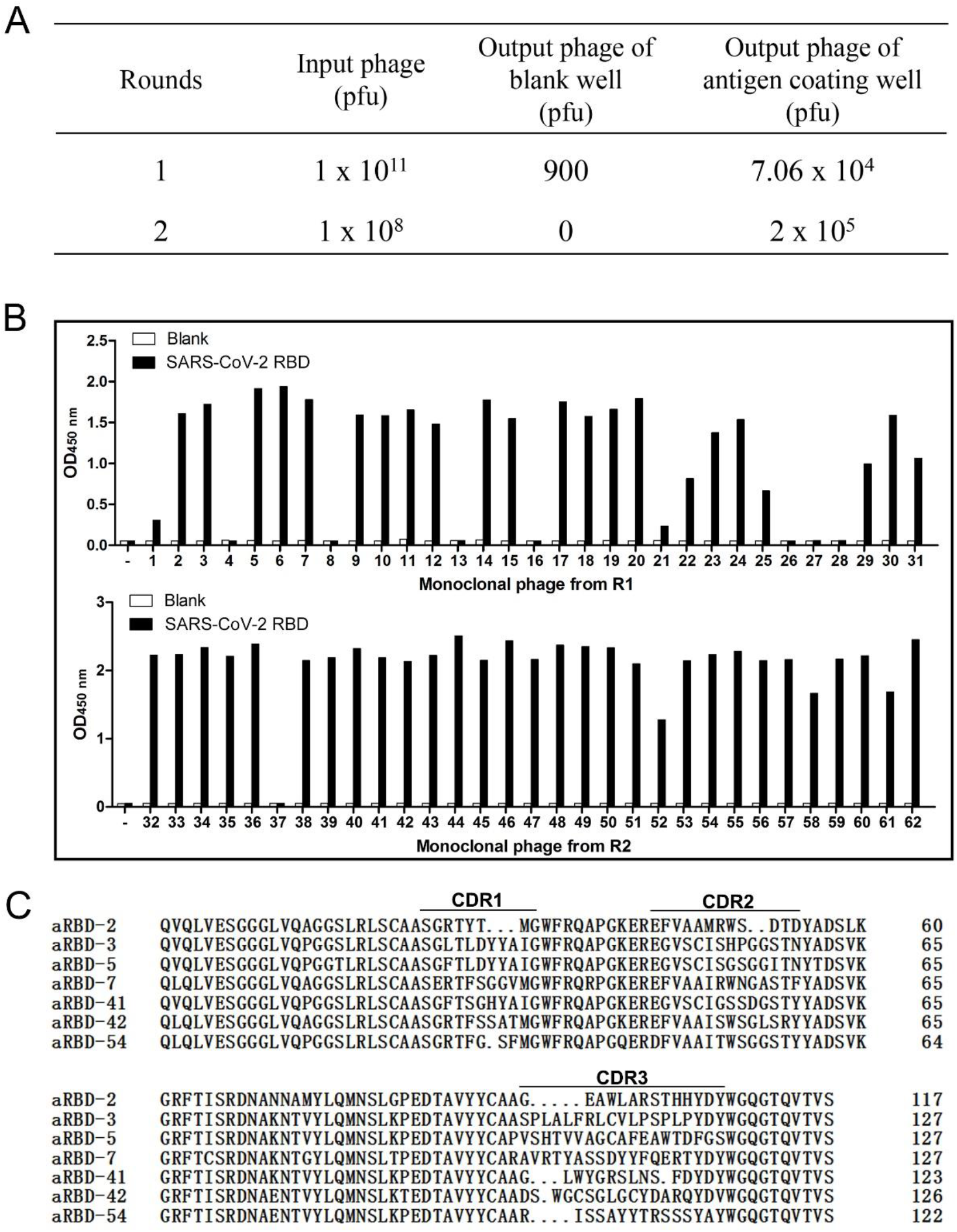
Isolation of anti-SARS-CoV-2 RBD Nbs from an immunized phage display library. (A) Enrichment results after panning on SARS-CoV-2 RBD. (B) Results of monoclonal phage ELISA. Randomly picked 31 individual clones from the 1st round of panning and 2nd round of panning were monitored against SARS-CoV-2 RBD and negative control (PBS). (C) Amino acid sequence of the isolated seven anti-RBD Nbs.

After each round of panning, thirty-one individual phages were randomly picked and their RBD-binding activity evaluated with phage ELISA. Nineteen and thirty phages were found to be positive for RBD binding after the first and second round of panning, respectively (**Fig. 1B**). Sequencing of the positive phage clones after two rounds of panning revealed seven unique Nbs (**Fig. 1C**), which were named as aRBD-2, aRBD-3, aRBD-5, aRBD-7, aRBD-41, aRBD-42 and aRBD-54. All seven phages can bind to the S1 domain of SARS-CoV-2 in ELISA, and one (aRBD-41) can also bind to SARS-CoV-1 RBD (**Fig. S2**).

### Binding characteristics of the identified nanobodies

The identified Nbs were expressed with a mammalian expression vector in 293F cells. To configure the Nb into IgG-like molecule, we fused the C-terminus of the identified Nbs to a TEV protease cleavage site and a human IgG1 Fc in a mammalian expression vector. The homo-bivalent Nb-TEV-Fc fusions were purified from the culture supernatant using protein A (**Fig. S3A**). All of the Nb-TEV-Fc fusions showed more than 100 mg/L yield after three days of expression (data not shown). To prepare Nb monomers without the Fc, the fusion proteins were digested with the TEV enzyme (6His tagged) and passed through protein G and Ni NTA column. Highly purified Nbs were obtained from the flow-through (**Fig. S3B**). The conformational stability of the seven Nbs were tested using circular dichroism, and the results showed that they were highly stable in solution, with the melting temperature exceeding 70°C (**Fig. S4**).

The SARS-CoV-2 RBD-binding abilities of the seven Nbs were first verified using size-exclusion chromatography (SEC). All seven Nbs formed stable complexes with RBD in solution (**Fig. 2A-I**). Furthermore, most Nb-Fc fusions demonstrated strong binding to both RBD and the entire ectodomain (S1+S2) of SARS-CoV-2 spike in ELISA, with EC_50_ of low nM. Compared to the human ACE2-Fc recombinant protein, they bind to the RBD with a higher affinity (**Fig. 2J**), while all but aRBD-42 bind to the entire ectodomain of spike protein with a higher affinity (**Fig. 2K**). In addition, we also tested the binding ability between the 7 Nbs and a RBD variant that contains N501Y point mutation derived from a recent new SARS-CoV-2 lineage that was rapidly spreading in UK [31]. As expected, N501Y variant showed an enhanced binding activity with ACE2-Fc than original RBD. Interestingly, all of the 7 Nbs exhibited similar binding activity to the variant and original RBD (**Fig. S5**).

**Fig. 2.**
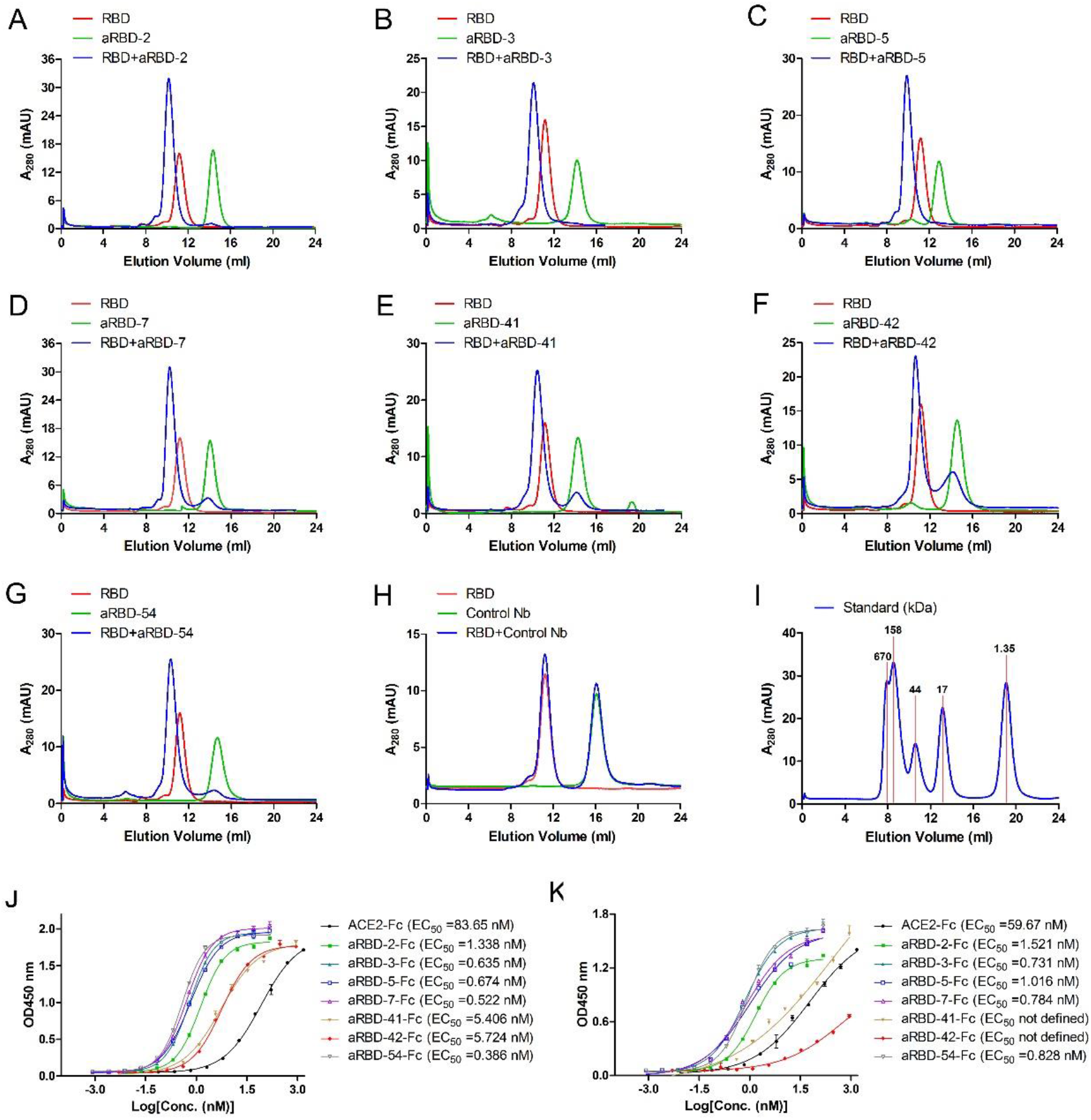
Size-exclusion chromatography (SEC) and ELISA analysis of interaction between SARS-CoV-2 RBD and Nbs in solution. SARS-CoV-2 RBD, Nbs and their 1:1 molar mixture were loaded over a Superdex 75 column (GE healthcare), respectively. (A)-(G) is the analysis curve of the seven Nbs, respectively; (H) is the curve of a negative control Nb; (I) is the curve of standard. If the Nbs could bind RBD to form complex, elution peak will move forward. SARS-CoV-2 RBD (J) and spike entire ectodomain (K) binding abilities of the purified Nb-Fc fusions were characterized using ELISA. EC_50_ was calculated by fitting the OD_450_ from serially diluted antibody with a sigmoidal dose-response curve. Error bars indicate mean ±SD from two independent experiments.

The binding affinity of the Nbs to RBD were also measured using Surface Plasmon Resonance (SPR). Six Nbs showed a high binding affinity, with K_D_ values of 2.60, 3.33, 16.3, 3.31, 21.9 and 5.49 nM for aRBD-2, aRBD-3, aRBD-5, aRBD-7, aRBD-41 and aRBD-54, respectively (**Fig. 3A-E and G**). Consistent with ELISA, aRBD-42 had a relatively weak binding affinity with a K_D_ of 113 nM (**Fig. 3F**). The affinities of Nb-Fc fusions were also measured by SPR. Probably due to dimerization, they showed an enhanced binding capability, with K_D_values ranging from 4.49 nM to 72.7 pM (**Fig. S6**).

**Fig. 3.**
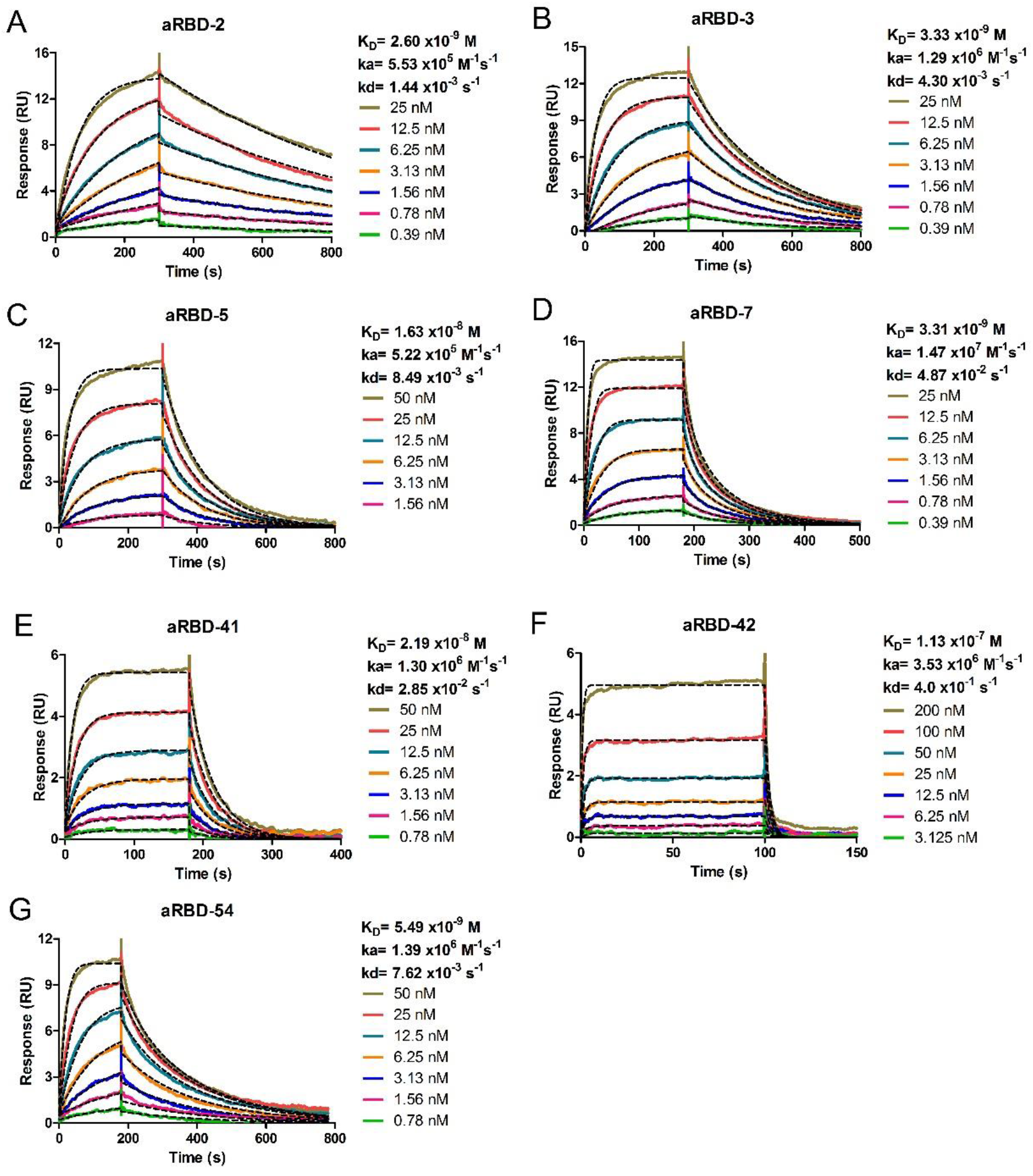
Characterization of binding affinity of isolated Nbs using SPR. Binding kinetics of aRBD-2 (A), aRBD-3 (B), aRBD-5 (C), aRBD-7 (D), aRBD-41 (E), aRBD-42 (F) and aRBD-54 (G) was measured by SPR, respectively. The SARS-CoV-2 RBD was immobilized onto a CM5 sensor chip. Nbs with serially 1:1 dilutions were injected and monitored by Biacore T200 system. The actual responses (colored lines) and the data fitted to a 1:1 binding model (black dotted lines) are shown.

### Nbs block RBD-ACE2 interaction

SARS-CoV-2 infection is initiated by the interaction of RBD and ACE2. To assess the ability of the Nbs in blocking RBD-ACE2 interaction, we performed competitive ELISA. Except for aRBD-42, which has the lowest RBD-binding affinity, all other Nbs (**Fig. 4A**) and their Fc fusions (**Fig. 4B**) effectively blocked the binding between ACE2-Fc and RBD in a dose dependent manner. Compare to monovalent Nbs, Nb-Fc fusions showed enhanced blocking activities with 5 to 90-fold decrease in half-maximal inhibitory concentration (IC_50_). The Nb-Fc fusions inhibited the binding of 10 nM ACE2-Fc to RBD with IC_50_ values at nanomolar level, consistent with their binding affinities.

**Fig. 4.**
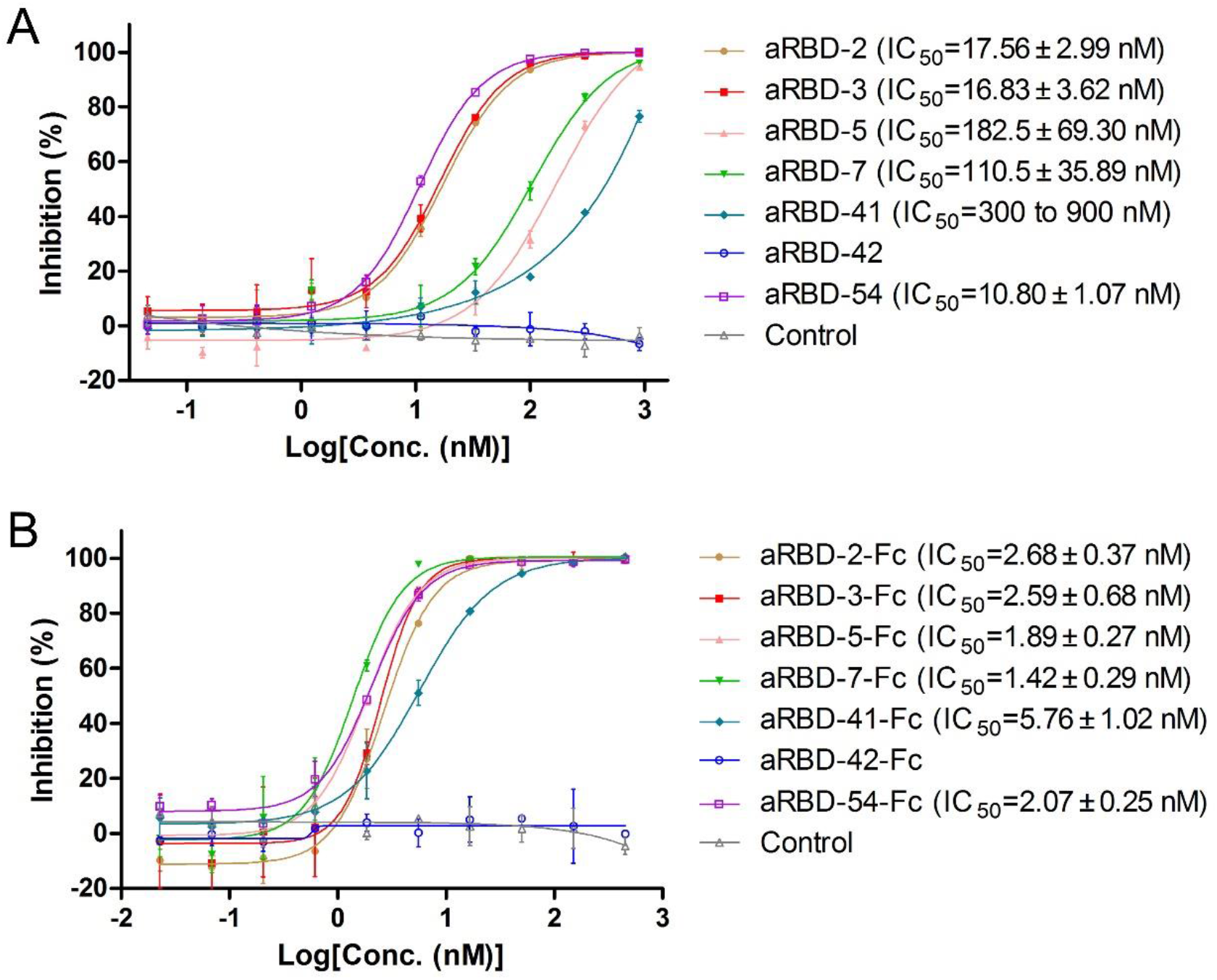
RBD-ACE2 blocking activities of isolated Nbs and their Fc fusions characterized with competitive ELISA. Competitive ELISA of ACE-Fc binding to SARS-CoV-2 RBD immobilized on the plates by increasing concentrations of Nbs (A) or Nb-Fc fusions(B). After the competition, bound ACE2-Fc (A) or biotinylated ACE2-Fc (B) was detected by HRP-anti-IgG1 Fc antibody or streptavidin-HRP, respectively. Error bars indicate mean ± SD from two independent experiments. IC_50 ,_was calculated by fitting the inhibition from serially diluted antibody to a sigmoidal dose-response curve.

### High affinity hetero-bivalent antibodies constructed depend on epitope grouping

To find out whether the Nbs bind to overlapping epitopes, the ability of the Nbs to compete with each other for ACE2 binding was studied with ELISA. The Nbs were serially diluted (ranging from 2.5 to 10240 nM) and used to compete with 5 nM of a Nb-TEV-Fc fusion to bind SARS-CoV-2 RBD coated on plates (**Fig. 5A-F**). The competition was summarized in **Fig. 5G**. Based on these grouping and an additional SEC results (**Fig. 6A and B**), we engineered two hetero-bivalent Nbs, namely aRBD-2-5 and aRBD-2-7, by connecting aRBD-2 head-to-tail with aRBD-5 and aRBD-7 through a (GGGGS)_3_ flexible linker, respectively. They were also expressed in 293F cells and purified as above (**Fig. 6C**). SEC indicated aRBD-2-5 and aRBD-2-7 were monomeric in solution (**Fig. 6D and E**), and circular dichroism spectrum analysis showed they were also highly stable in solution (**Fig. S4h, i**).

**Fig. 5.**
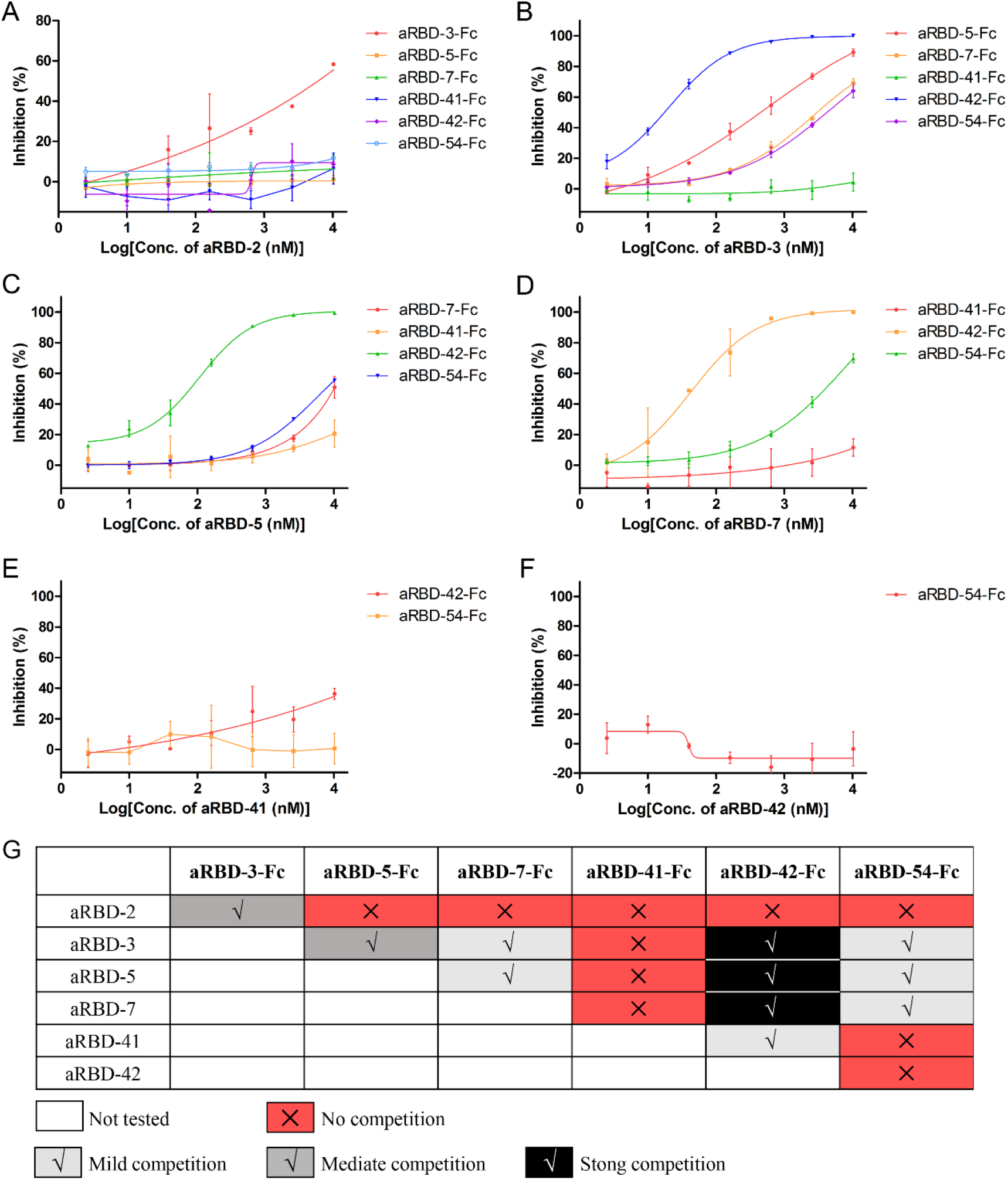
Epitope grouping results of the seven identified nanobodies. aRBD-2 (A), aRBD-3 (B), aRBD-5 (C), aRBD-7 (D), aRBD-41 (E) and aRBD-42 (F) was competed with other Nb-Fc fusions to bind SARS-CoV-2 RBD immobilized on the plates. Competition was determined by the reduction of HRP-anti-IgG1 Fc induced chemiluminescence signal (OD_450 nm_). The inhibition was calculated by comparing to the Nb negative control well. Error bars indicate mean ± SD fro m two independent experiments. Competition strength is negatively correlated with OD_450 nm_ signal, the strength was summarized (G). aRBD-2 and aRBD-41 only showed moderate and mild competition with aRBD-3 and aRBD-42 for RBD binding, respectively. aRBD-3, aRBD-5, aRBD-7 and aRBD-42 showed mild to strong competition with each other for RBD binding. aRBD-3, aRBD-5, aRBD-7 and aRBD-54 also showed mild to strong competition with each other for RBD binding. aRBD-42 had no competition with aRBD-54.

**Fig. 6.**
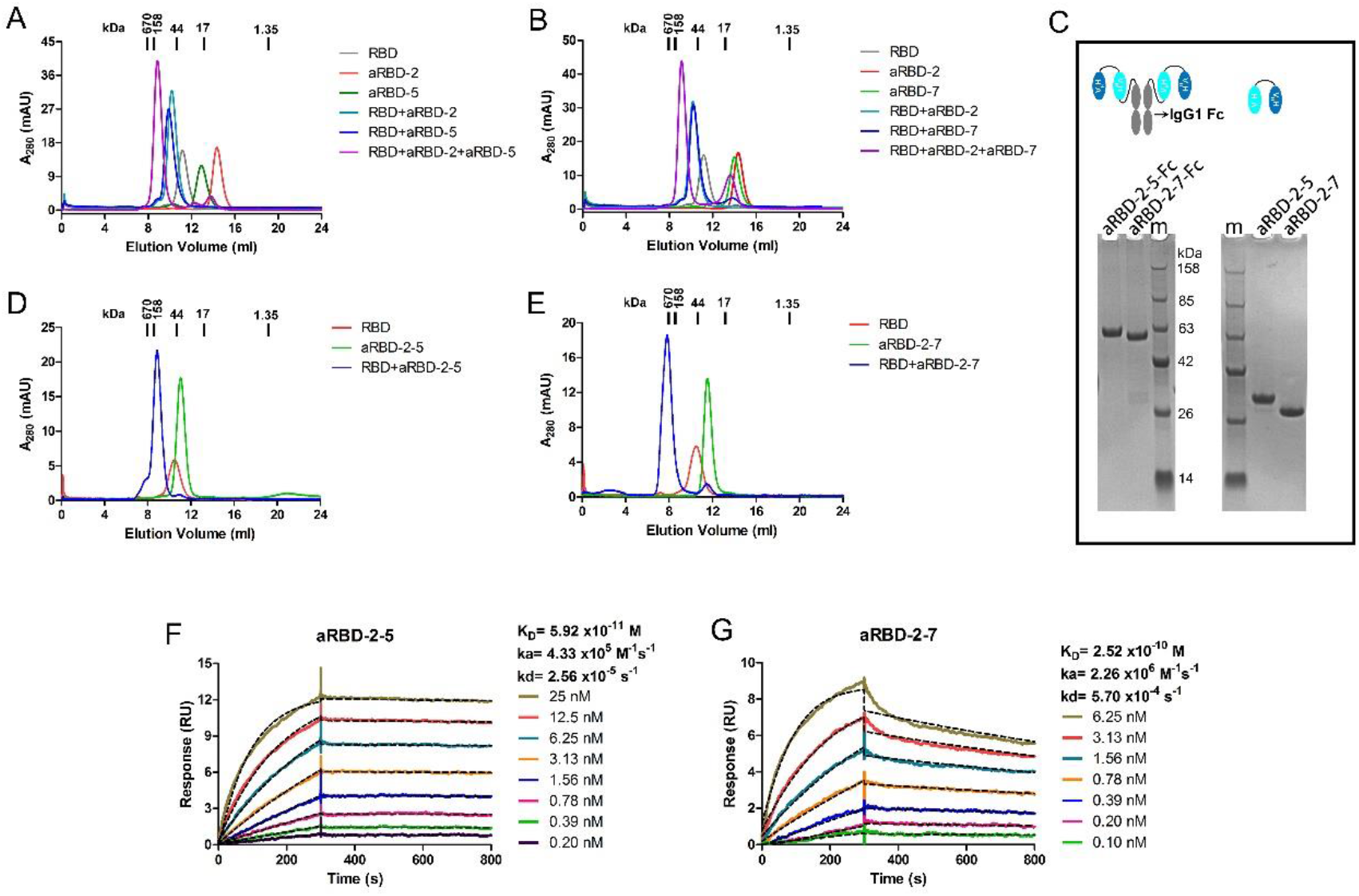
Preparation and characterization of the hetero-bivalent Nbs. (A) SEC profiling of RBD, aRBD-2 and aRBD-5 complex. (B) SEC profiling of RBD, aRBD-2 and aRBD-7 complex. (C) Reduced SDS-PAGE results of purified aRBD-2-5, aRBD-2-7 and their Fc fusions. (D) SEC profiling of RBD and aRBD-2-5 complex. (E) SEC profiling of RBD and aRBD-2-7 complex. The SARS-CoV-2 RBD binding kinetics of aRBD-2-5 (F) and aRBD-2-7 (G) was measured by SPR, respectively. The two hetero-bivalent Nbs with serially 1:1 dilutions were injected and monitored by Biacore T200 system. The actual responses (colored lines) and the data fitted to a 1:1 binding model (black dotted lines) are shown.

The RBD binding activities of aRBD-2-5 and aRBD-2-7 were studied with SEC (**Fig. 6D and E**) and SPR. In contrast to the monovalent Nbs, the hetero-bivalent aRBD-2-5 and aRBD-2-7 showed a greatly enhanced binding affinity, with K_D_ values of 59.2 pM and 0.25 nM, respectively (**Fig. 6F and G**). Similarly, their Fc fusions also showed an enhanced binding affinity, with K_D_ values of 12.3 pM and 0.22 nM, respectively (**Fig. S6H and I**).

### Hetero-bivalent Nbs exhibit potent neutralizing ability against live SARS-CoV-2

To assess the ability of the Nbs in neutralizing SARS-CoV-2, we developed a SARS-CoV-2 micro-neutralization assay and assessed representative Nbs in this assay. Nbs that were serially diluted to different concentrations were incubated with ∼ 200 PFU of SARS-CoV-2 and inoculated onto Vero E6 cells in 96-well plates. The inoculum was removed after 1 hour, and the cells were covered with semi-solid medium for 2 days, before the infection was assessed by an immunofluorescence assay utilizing antibodies specific for SARS-CoV-2 N protein. Three representative monomeric Nbs, aRBD-2, aRBD-5, and aRBD-7, showed only modest level of neutralization at antibody concentrations of 33 to 100 μg/ml (**Fig. S7A-C**). aRBD-2 was more effective than aRBD-5 and aRBD-7 in neutralizing SARS-CoV-2, correlating with its higher binding affinity to RBD. By contrast, the dimeric Nbs showed greatly enhanced neutralizing potency. The homo-bivalent aRBD-2-Fc, aRBD-5-Fc and aRBD-7-Fc exhibited 50% neutralization dose (ND_50_) of 0.092 μg/mL (∼1.12 nM), 0.440 μg/mL (∼5.34 nM) and 0.671 μg/mL (∼8.02 nM), respectively (**Fig. S8A-C and Fig. 7E**), again correlating with their RBD binding affinities. Interestingly, the hetero-bivalent Nbs exhibited an even higher neutralizing potency than the homo-dimeric Nbs. The fitted ND_50_ for aRBD-2-5 and aRBD-2-7 is 1.22 ng/mL (∼0.043 nM) and 3.18 ng/mL (∼0.111 nM), respectively (**Fig. 7A, B and E**). The Fc fusions of the hetero-bivalent Nbs did not further increase the neutralization potency. The ND_50_ for aRBD-2-5-Fc and aRBD-2-7-Fc is 11.8 ng/mL (∼0.107 nM) and 6.76 ng/mL (∼0.0606 nM), respectively (**Fig. 7C, D and E**).

**Fig. 7.**
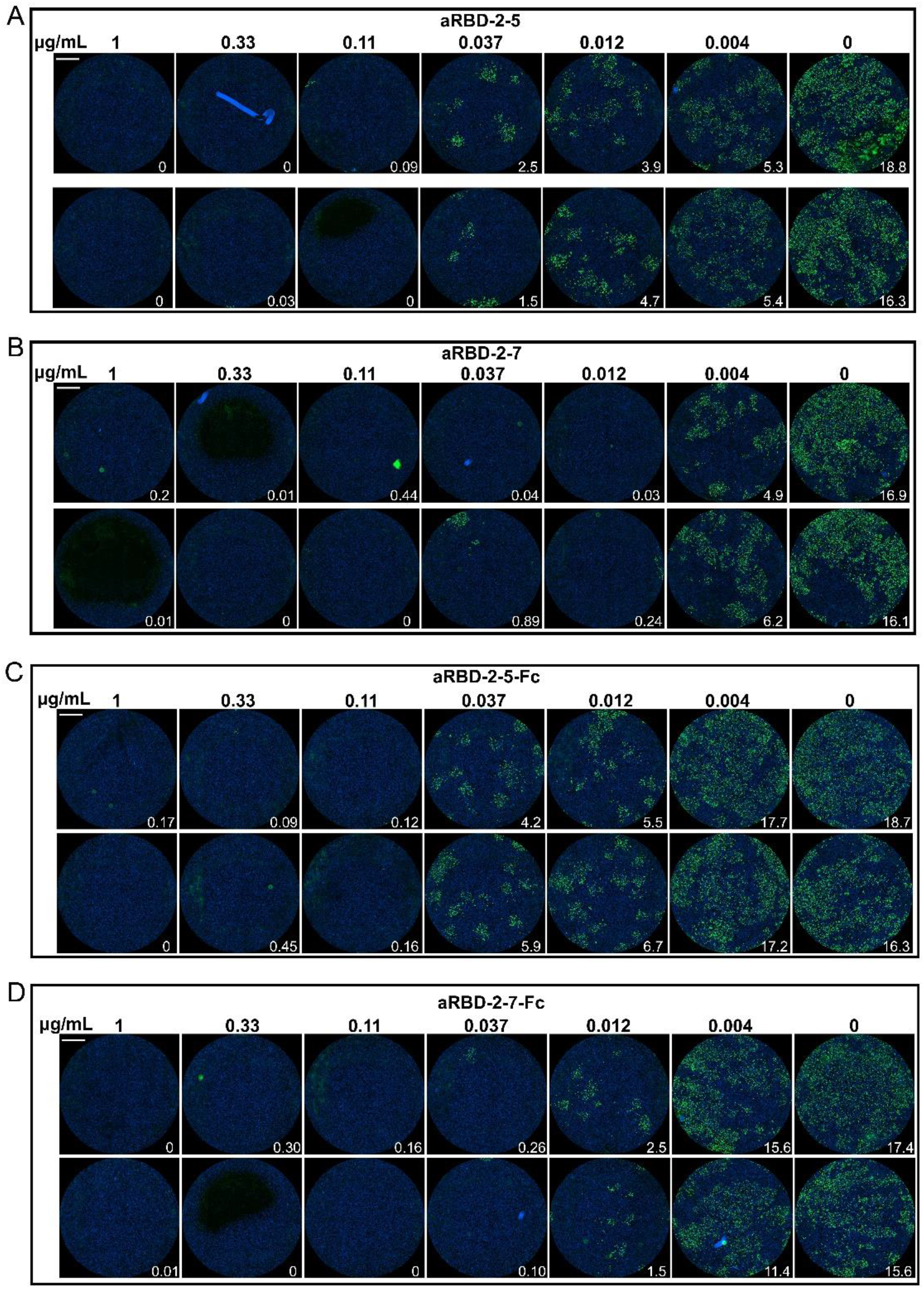
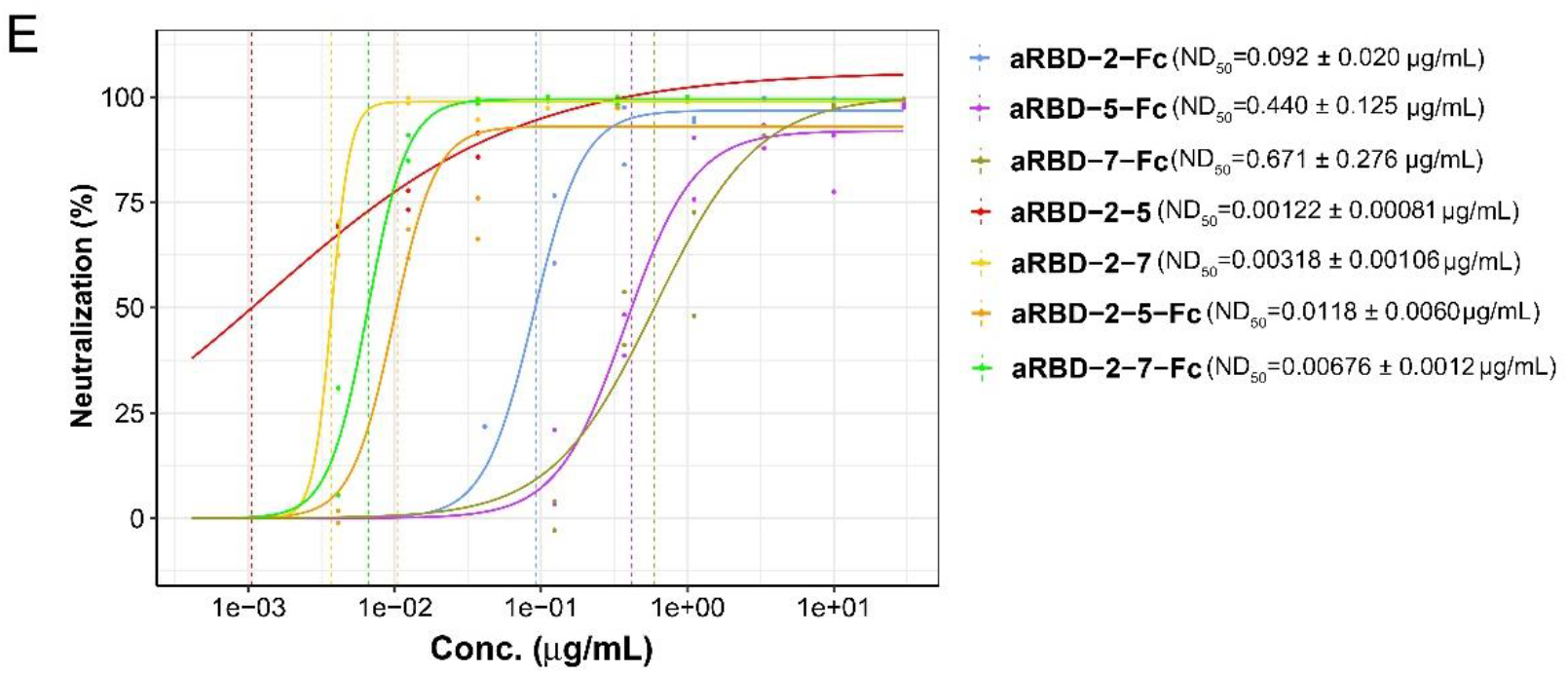
In vitro SARS-CoV-2 neutralization of the hetero-bivalent Nbs and their Fc fusions. The serially diluted aRBD-2-5 (A), aRBD-2-7 (B), aRBD-2-5-Fc (C) and aRBD-2-7-Fc (D) was incubated with ∼200 PFU of SARS-CoV-2. The mixture were then added to Vero E6 cells in 96-well plates. After 2 days of infection, the infected virus were stained green with a monoclonal antibody against SARS-CoV-2 NP and an Alexa Fluor 488-conjugated goat anti-mouse secondary antibody. The nucleus was stained blue with Hoechst 33342. Each experiment was performed in duplicate. Numbers in the low right corner of each grid were the percentage of cells infected by the virus. (E) ND_50_ of the identified Nbs and their Fc fusions were calculated by fitting the neutralization from serially diluted antibody with a Log-logistic model. The data are fitted to the model with the drc package in R to obtain the 95% confidence intervals.

## Discussion

The infection of epithelial cells by SARS-CoV-2 is initiated by the interaction between the Spike RBD and ACE2 [25, 32]. Hence, RBD-targeting antibodies hold promise as prophylactics and therapeutics for SARS-CoV-2. Here, seven unique Nbs were isolated from RBD-immunized alpacas and their CDR sequences are distinct from other reported Nbs [33-41]. Four of the Nbs exhibited high affinity of low-nanomolar K_D_ (**Fig. 3**), similar to recently reported natural [33, 34] and synthetic Nbs [35-37]. We also measured the RBD-binding affinity of the Nb-Fc fusions. Due to the bivalent nature and ∼6-fold increase in the molecular weight, most of the Nb-Fc chimeric antibodies showed a higher affinity (K_D_ ranging from 72.7 pM to 4.5 nM) than their monomeric counterparts (**Fig. S6**). This affinity is even higher than that of some monoclonal antibodies isolated from lymphocytes of convalescent COVID-19 patients [16- 18].

Except for the Nb with the lowest affinity for RBD (aRBD-42; K_D_ of 113 nM), the other six Nbs are all capable of blocking the interaction between ACE2 and RBD. aRBD-2, aRBD-3 and aRBD-54, which had a higher RBD-binding affinity, showed a stronger ACE-RBD blocking capacity than aRBD-5 and aRBD-41 (**Fig. 4A**). However, aRBD-7, which had a similarly high RBD binding affinity of 3.31 nM, only exhibited a weak ACE2-RBD blocking activity (**Fig. 4A**). We thus infer that different Nbs may occupy different epitopes on RBD, leading to varying strength of ACE2 binding interference. The epitopes of some Nbs may overlap more closely with that of ACE2. Interestingly, even when the Nbs with a relatively weak ACE2-RBD blocking ability were fused with IgG1 Fc to form homodimers, their blocking ability were increased more than 75-fold (**Fig. 4A and B**). This effect is probably due to the increased apparent RBD-binding affinity by dimerization as well as the additional steric hindrance caused by the increased size. Further investigations are needed to understand the underlying mechanisms.

According to grouping results of the seven Nbs, two hetero-bivalent antibodies were constructed by fusing aRBD-2 to aRBD-5 and aRBD-7 tail-to-head with a flexible linker, which achieved a more than 10-fold increase in RBD-binding affinity (**Fig. 6F and G**). Consistent with the increased affinity and steric hindrance, the SARS-CoV-2 neutralization potency of aRBD-2-5 and aRBD-2-7 were greatly enhanced, with ND_50_ of 1.2 ng/mL (∼0.043 nM) and 3.2 ng/mL (∼0.111 nM) (**Fig. 7**).The neutralization potency of our aRBD-2-5 and aRBD-2-7 appears to be better than other previously reported Nbs, their engineered form [33-40], and some traditional human monoclonal antibodies[13-18]. However, they appear to be less potent than a recently reported multivalent Nb[41].

In summary, we have identified several high-affinity natural Nbs with RBD-ACE2 blocking ability and two hetero-bivalent Nbs with potent SARS-CoV-2 neutralization capacity. Alpaca V_H_H has a high degree of homology with human V_H_3, so it has low immunogenicity in humans [42, 43]. These Nbs can be further improved with respect to their antiviral function through affinity maturation or genetic modification, potentially serving as therapeutics for treating COVID-19.

## Material and Methods

### Protein expression and purification

The coding sequences for SARS-CoV-2 RBD (aa 321-591), SARS-CoV-2 RBD (aa 321-591, N501Y), SARS-CoV-2 S1 (aa 1-681), SARS-CoV-1 RBD (aa 309-540), human ACE2 extracellular domain (aa 19-615) and the identified Nbs, were appended with a TEV enzyme site and a human IgG1 Fc at the C-terminus as well as the IFNA1 signal peptide at the N-terminus. The fusions were cloned into the mammalian expression vector pTT5. The expression vectors were transiently transfected to human HEK293F cells with polyethylenimine (Polyscience). Three days later, cell supernatants were obtained by centrifugation at 3000 g for 10 min, diluted 1:1 with the running buffer (20 mM Na_2_HPO_4_, 150 mM NaCl, pH 7.0), and loaded on protein A column. The bound protein was eluted with 100 mM acetic acid on ÄKTA pure (GE healthcare). To remove IgG1 Fc, the purified fusion proteins were first digested with 6xHis-tagged TEV enzyme. Protein A (Protein G for nanobodies) and Ni NTA were then used sequentially to remove the undigested fusion protein, Fc and the TEV enzyme. Fc-free recombinant proteins were collected from the flow-through. Protein purity was estimated by SDS-PAGE (**Fig. S1**), and the concentration was measured using the spectrophotometer (analytikjena).

### Phage display library construction

The experiments involved alpacas were approved by a local ethics committee. Two female alpacas were immunized by 2 times of subcutaneous injection and 1 time of intramuscular injection, each with 500 μg SARS-CoV-2 RBD in PBS, which was emulsified with an equal volume of Freund’s adjuvant (Sigma Aldrich). Two weeks after the final boost, more than 1 × 10^7^ lymphocytes were isolated from peripheral blood by Ficoll 1.077 (Sigma Aldrich) separation, and the total RNA from the lymphocytes was isolated using Total RNA kit (omegabiotek) according to the manufacturer’s protocol. First strand cDNA synthesis was performed with 4 μg of total RNA per reaction using PrimeScript^™^ II 1st Strand cDNA Synthesis Kit and oligo-dT primer (TAKARA) according to the manufacturer’s protocol. The variable domain of heavy-chain only antibody (V_H_H) was amplified by PCR using the following primers (Forward primer: GCTGCACAGCCTGCTATGGCACAGKTGCAGCTCGTGGAGTCTGGGGG; Reverse primer: GAGTTTTTGTTCGGCTGCTGCTGAGGAGACGGTGACCTGGGTCCCC). The phagemid pR2 was amplified by PCR using the following primers (Forward primer: AGCAGCCGAACAAAAACTCATCTCAGAAGAG; Reverse primer: CCATAGCAGGCTGTGCAGCATAGAAAGGTACCACTAAAGGAATTGC). Two pmol of the V_H_H fragments and 0.5 pmol of the amplified pR2 vector were mixed and diluted to 50 μl. An equal volume of 2x Gibson Assembly mix was added to the mixture and incubated at 50°C for 1 hour. The ligation was cleaned up by Cycle Pure Kit (omegabiotek) and transformed into TG1 electro-competent cells in 0.1 cm electroporation cuvette using BTX ECM 399 Electroporation System (Harvard Apparatus) with the following setting: 2.5 kV, 5 ms. The transformants were spread on five 150 mm TYE agar plates supplemented with 2% glucose and 100 μg/mL ampicillin, followed by overnight culturing at 37 °C. The colonies were scraped from the plates with a total of 20 mL 2×TY medium and thoroughly mixed. 200 μL of the liquid was inoculated to 200 mL 2×TY to amplify the library. Phage particles displaying V_H_H were rescued from the library using KM13 helper phage.

### Biopanning and selection of positive clones

Two rounds of panning were performed. Immuno MaxiSorb plates (Nunc) were coated with 0.1 mL of SARS-CoV-2 RBD solution (100 and 20 μg/mL in the 1st and 2nd round, respectively). Control wells without antigen coating were used in parallel in every round of panning. After blocking with MPBS (PBS supplemented with 5% milk powder) for 2 h at room temperature (RT), 1×10 ^11^ pfu of the library phages were added for the 1st round of selection. The wells were washed with PBST (PBS supplemented with 0.1% tween-20) for 20 times to remove the unbound phages. Bound phages were eluted by digestion with 100 μL of 0.5 mg/mL trypsin for 1 h at RT. The eluted phages were used to infect E. coli TG1 for titer determination and amplification. The 2nd round of panning was performed similarly with the following differences: the amount of input phage was 1×10 ^8^ pfu, the washing time was 30 times, and the concentration of tween-20 in washing buffer was 0.2%.

Thirty-one individual clones from each round of panning were picked and identified using monoclonal phage ELISA. The monoclonal phage was rescued with helper phage KM13 and added to the well coated with 0.1 μg of RBD. After 1 h of incubation at RT, the wells were washed 4 times with PBST and added with HRP-anti-M13 antibody. After washing 4 times with PBST, TMB (Beyotime) was added to each well and incubated in the dark at RT for 2 min. The chromogenic reaction was stopped with 50 μL of 1 M sulfuric acid, and OD_450 nm_ was determined. The clone with OD_450 nm_ that was 20 times higher than that of the control well is defined as a positive clone. The phagemids extracted from the positive clones were sequenced.

### Size-exclusion chromatography

The interaction of SARS-CoV-2 RBD and the Nbs in solution was studied with gel filtration. SARS-CoV-2 RBD, Nbs and their mixture (1.6 nmol of SARS-CoV-2 RBD mixed with 1.6 nmol of Nbs) were run over a Superdex 75 column (GE healthcare) at a flow rate of 0.5 mL/min with AKTA pure.

### Enzyme-linked immunosorbent assay (ELISA)

Immuno MaxiSorb plates (Nunc) were coated and blocked as above. For non-competitive ELISA of purified Nb-Fc and ACE2-Fc binding assay, Nb-Fc and ACE2-Fc solutions that were serially diluted 1:3 were added to the plates and incubated for 1 h at RT. After washing with PBST 4 times, the bound Nb-Fc and ACE2-Fc were detected with a monoclonal anti-IgG1 Fc-HRP antibody (sino Biological). For characterizing the epitope competition between the identified Nbs, serially 1:4 diluted Nb solutions (ranging from 2.5 to 10240 nM) were mixed with 5 nM of Nb-Fc solutions. After incubation in RBD coated wells and standard washing, bound Nb-Fc was detected with a monoclonal anti-IgG1 Fc-HRP antibody. For ACE2-RBD blocking assay, serially 1:3 diluted Nb solutions (ranging from 0.046 to 900 nM) and Nb-Fc solution (ranging from 0.023 to 450 nM) were mixed with 10 nM of ACE2-Fc and 10 nM of biotinylated ACE2-Fc, respectively. After incubation in RBD coated wells and standard washing, bound ACE2-Fc and biotinylated ACE2-Fc was detected with an anti-IgG1 Fc-HRP antibody or HRP-streptavidin, respectively. The chromogenic reaction and OD_450 nm_ measurement were performed similarly as described for phage ELISA.

### Circular Dichroism (CD)

Secondary structure and thermal stabilities of identified Nbs were studied by CD spectra using a Chirascan Spectrometer (Applied Photophysics). Prior to CD measurements, the sample buffer was changed to phosphate-buffered saline (PBS), and the protein concentration was adjusted to 0.3 mg/ml. The CD spectra were acquired for each sample from 180 to 260 nm using a 1 mm path length cell. For thermal titration, CD spectra were acquired between 20°C to 95°C with temperature steps of 2.5°C. CD signals at 205 nm were used to characterize the structural changes during thermal titration. Each experiment was repeated twice, and the data were fitted with Prism to obtain the Tm values.

### Surface Plasmon Resonance (SPR)

SPR measurements were performed at 25°C using a BIAcore T200 system. SARS-CoV-2 RBD was diluted to a concentration of 15 μg/mL with sodium acetate (pH 4.5) and immobilized on a CM5 chip (GE Healthcare) at a level of ∼150 response units (RU). All proteins were exchanged into the running buffer (PBS (pH 7.4) supplemented with 0.05% Tween 20), and the flow rate was 30 μL/min. The blank channel of the chip served as the negative control. For affinity measurements, a series of different concentrations of antibodies were flowed over the sensorchip. After each cycle, the chip was regenerated with 50 mM NaOH buffer for 60-120 seconds. The sensorgrams were fitted with 1:1 binding model with the Biacore evaluation software.

### SARS-CoV-2 neutralization assay

Nbs and Nb-Fc fusions in a three-fold dilution concentration series were incubated with ∼200 plaque-forming units (PFU) of SARS-CoV-2 (USA-WA1/2020 isolate) for 30 minutes. The antibody and virus mixture were then added to Vero E6 cells in 96-well plates (Corning). After one hour, the supernatant was removed from the wells, and the cells were washed with PBS and overlaid with DMEM containing 0.5% methyl cellulose. After 2 days of infection, the cells were fixed with 4% paraformaldehyde, permeabilized with 0.1% Triton-100, blocked with DMEM containing 10% FBS, and stained with a rabbit monoclonal antibody against SARS-CoV-2 NP (GeneTex, GTX635679) and an Alexa Fluor 488-conjugated goat anti-mouse secondary antibody (ThermoFisher Scientific). Hoechst 33342 was added in the final step to counterstain the nuclei. Fluorescence images of the entire well were acquired with a 4x objective in a Cytation 5 (BioTek). The total number of cells, as indicated by the nuclei staining, and the infected cells, as indicated by the NP staining, were quantified with the cellular analysis module of the Gen5 software (BioTek). All experiments involving live SARS-CoV-2 were carried out under BSL-3 containment. A Log-logistic model[44] was used to model the dose-response curves of the antibodies. The data are fitted to the model with the drc package in R to obtain the 95% confidence intervals and ND_50_. It should be noted that all above Nbs were lyophilized at concentration of 2-5 mg/mL and kept at room temperature for one week for transportation. The lyophilized Nbs were re-dissolved in ddH_2_O before they were used in neutralization assay.

## Acknowledgements

We would like to thanks all the staff who participated in this work for their important contribution. This work is supported by the Strategic Priority Research Program of the Chinese Academy of Sciences (Grant No. XDB29030104), the National Natural Science Foundation of China (Grant No.: 31870731, 31971129 and U1732109), the 100 Talents Programme of The Chinese Academy of Sciences, the Fundamental Research Funds for the Central Universities (WK2070000108), the COVID-19 special task grant supported by Chinese Academy of Science Clinical Research Hospital (Hefei) (Grant No. YD2070002017), the Innovation team cultivation fund of USTC, Jack Ma Foundation and the New medical science fund of USTC (WK2070000130). The *in vitro* SARS-CoV-2 neutralization experiments were supported by a COVID-19 pilot grant from UTHSCSA to Y.X.

## Conflict of interest

All the authors have no conflicts of interest. Patents have been applied (No.: CN202011037351.1 and CN202011037426.6, Filing Date: Sep 25, 2020).

## Author Contributions

Tengchuan Jin and Yan Xiang provide funding, designed the study, participated in data analysis, and wrote the manuscript. Huan Ma and Weihong Zeng designed the study, performed the majority of experiments, analyzed the data and drafted the manuscript. Other authors participated in the experiments and/or writing of the manuscript.

---

Supplementary materials: Fig.S1 to Fig. S7.

Supplemental Figures

**Fig. S1.**
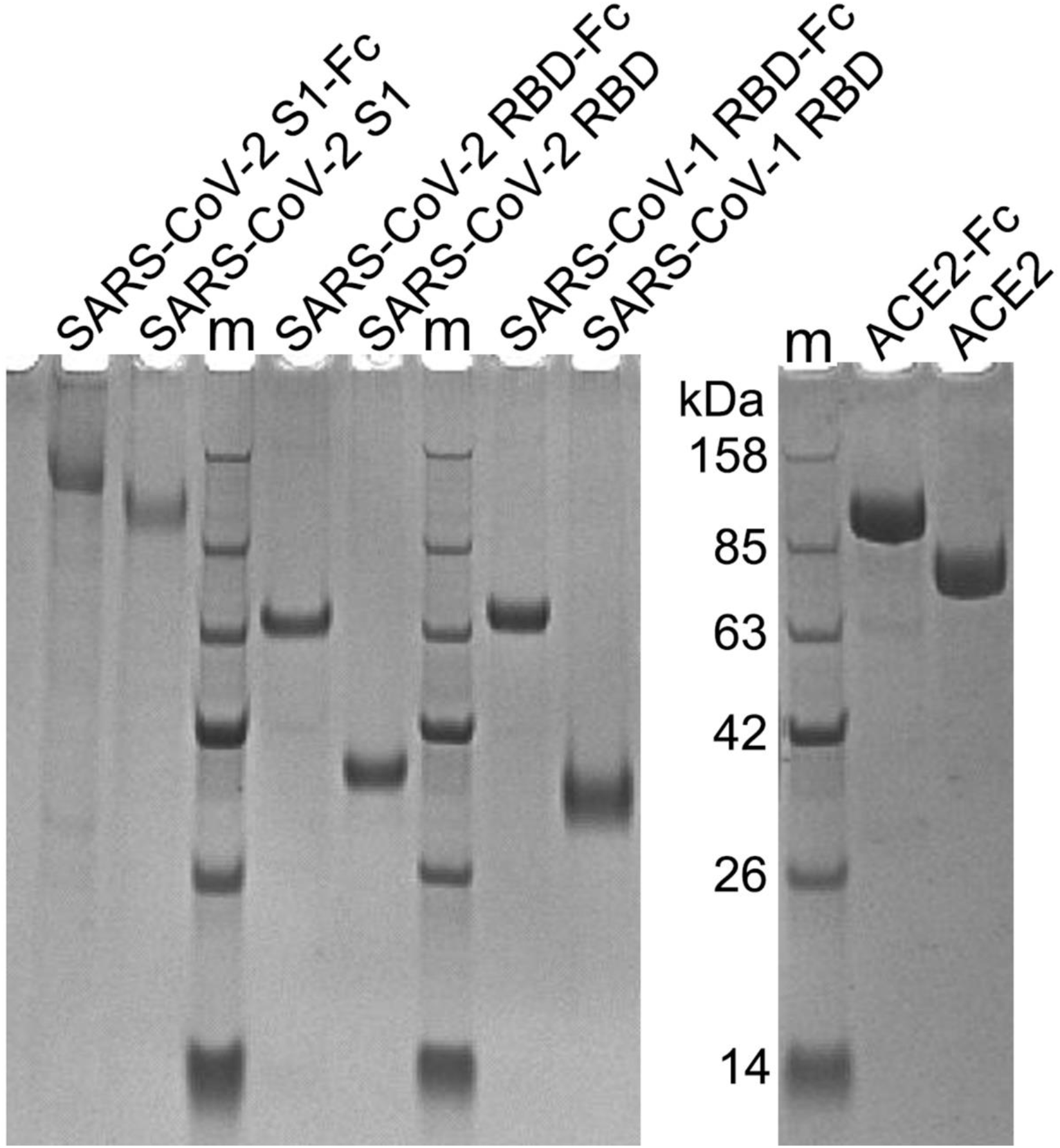
Reduced SDS-PAGE analysis of the purified proteins used in this study. All the proteins were fused with a TEV protease cleavage site and human IgG1 Fc and expressed using HEK293F cells, all the protein-TEV-Fc fusions were purified from culture supernatant with protein A. After digesting with TEV enzyme, the proteins without TEV-Fc were purified from the flow-through of protein A and Ni NTA. “m” is marker.

**Fig. S2.**
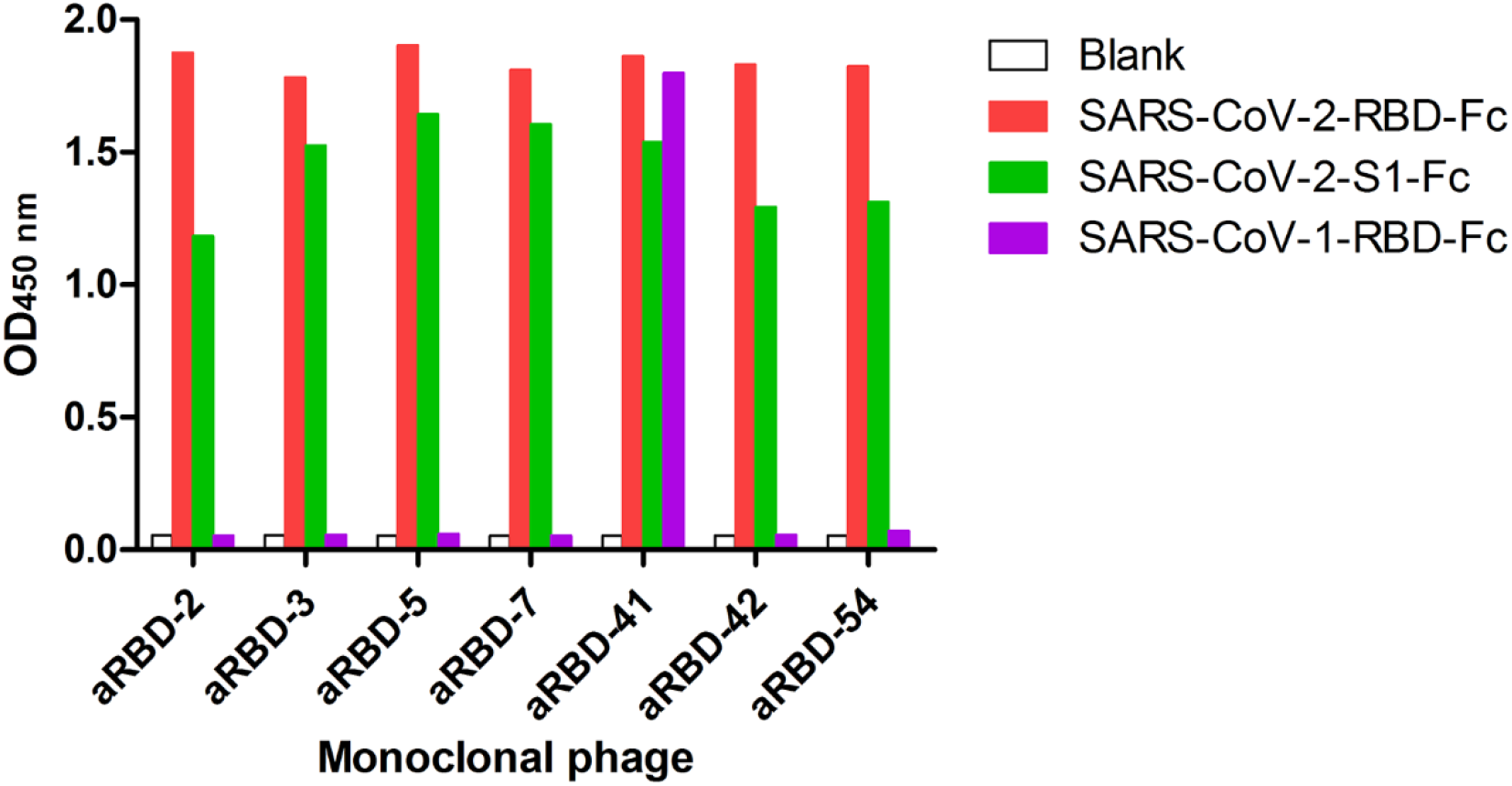
Identification of the binding of the seven positive phage clones to SARS-CoV related antigens using phage ELISA. All seven phages can bind to the S1 domain of SARS-CoV-2, one (aRBD-41) of them can also bind to SARS-CoV-1 RBD.

**Fig. S3.**
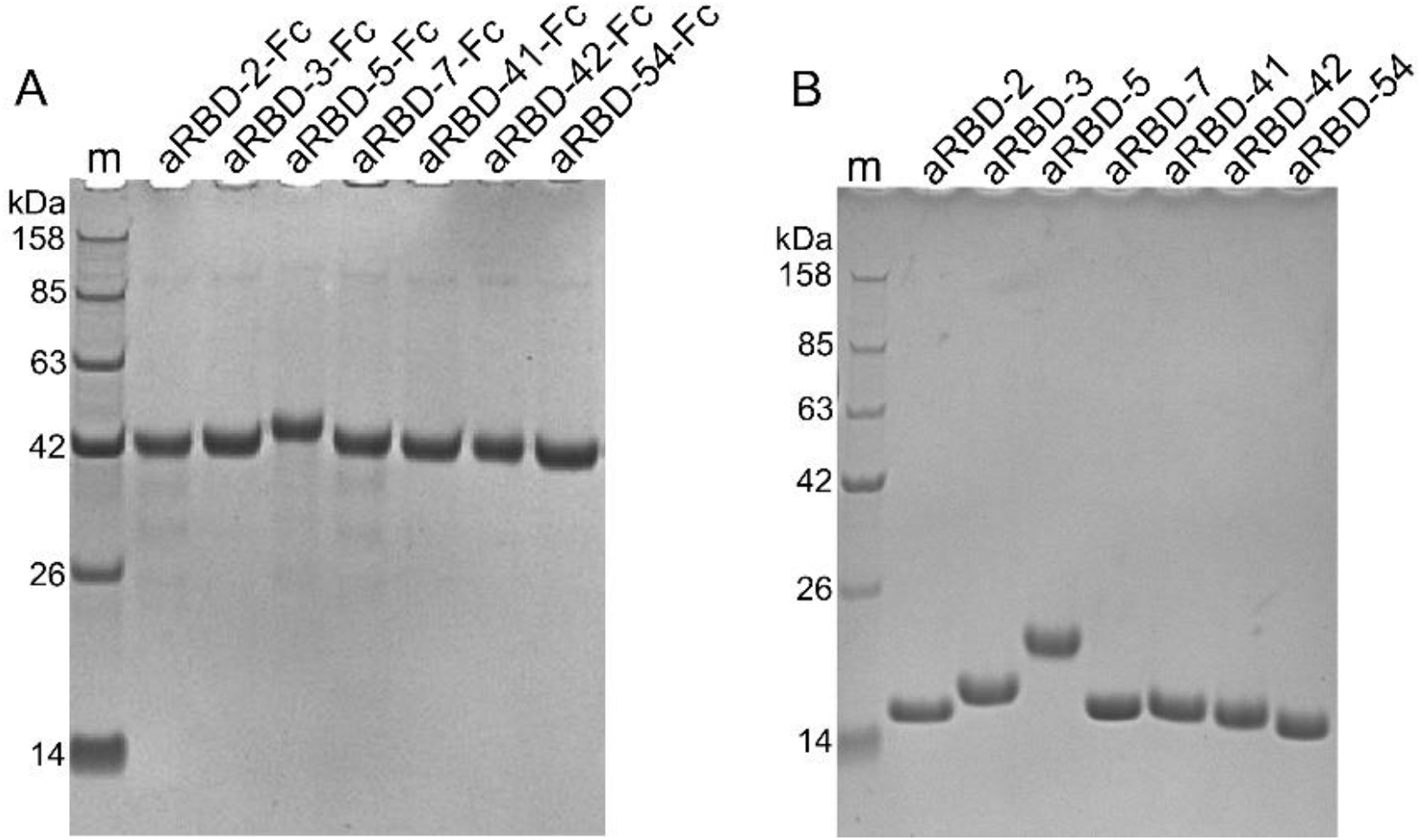
Reduced SDS-PAGE analysis of the purified Nbs and their Fc fusions. (A) The purified seven Nb-Fc fusions; (B) The purified seven Nbs. Due to glycosylation, the molecular weight of aRBD-5 is higher than others.

**Fig. S4.**
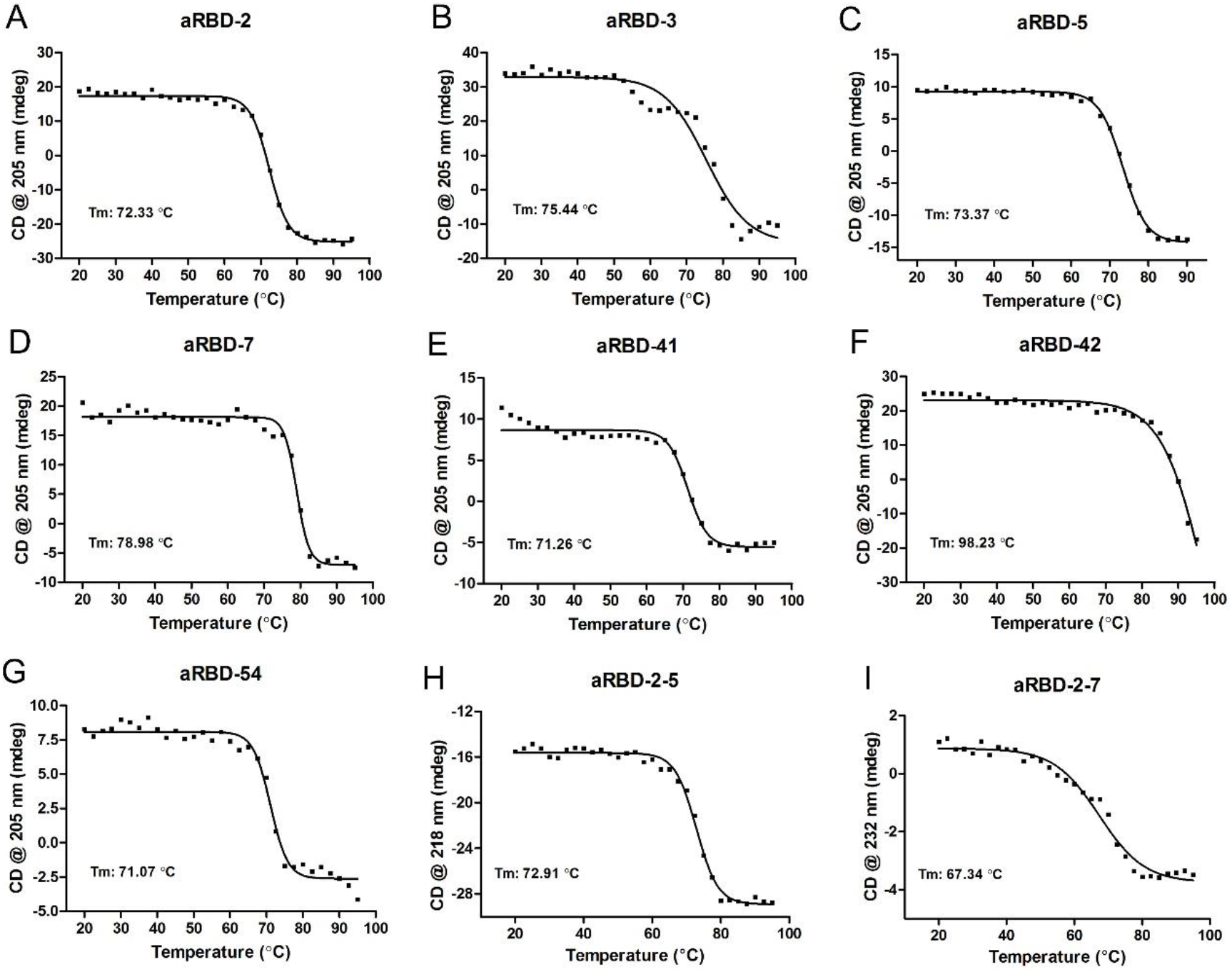
Thermal denature of Nbs by CD spectrum. A-I is thermal denature curve of aRBD-2, aRBD-3, aRBD-5, aRBD-7, aRBD-41, aRBD-42, aRBD-54, aRBD-2-5 and aRBD-2-7, respectively. Each experiment was repeated twice, the results data were fitted by Prism software.

**Fig. S5.**
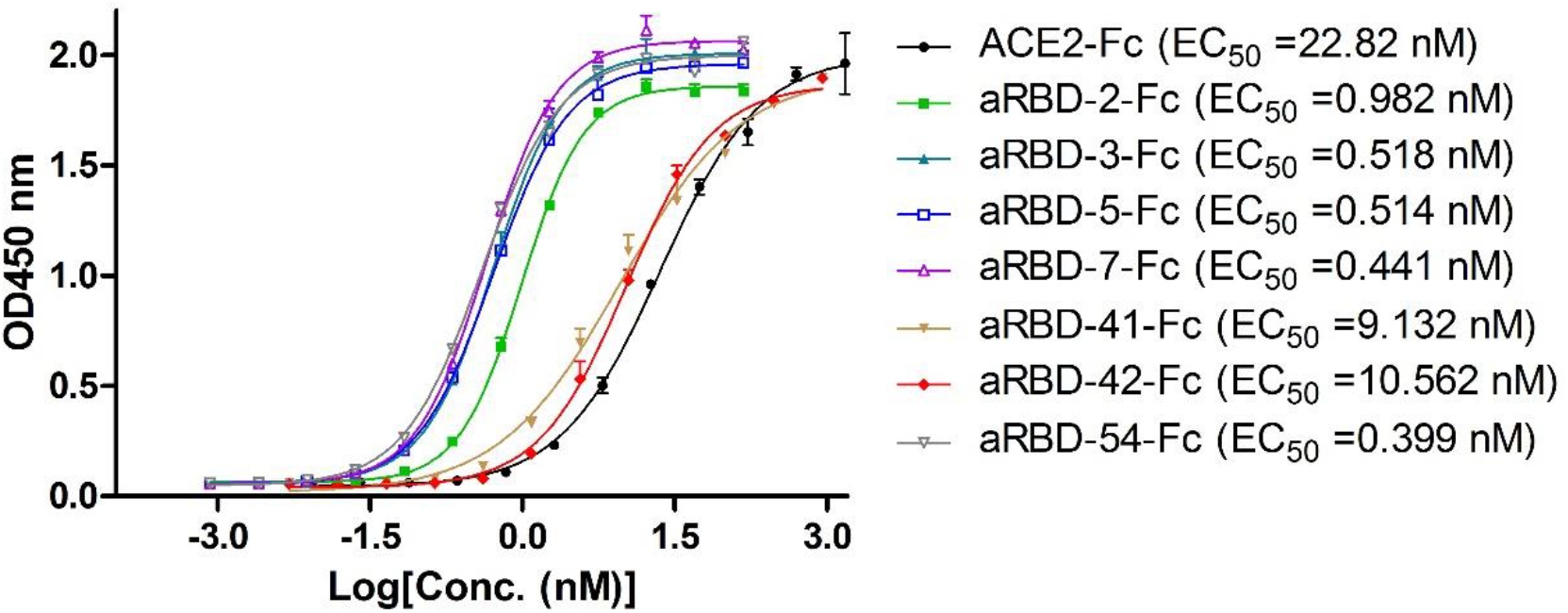
ELISA results for characterization of binding between identified Nbs and RBD variant contains N501Y mutation. EC_50_ was calculated by fitting the OD_450_ from serially diluted antibody with a sigmoidal dose-response curve. Error bars indicate mean ±SD from two independent experiments.

**Fig. S6.**
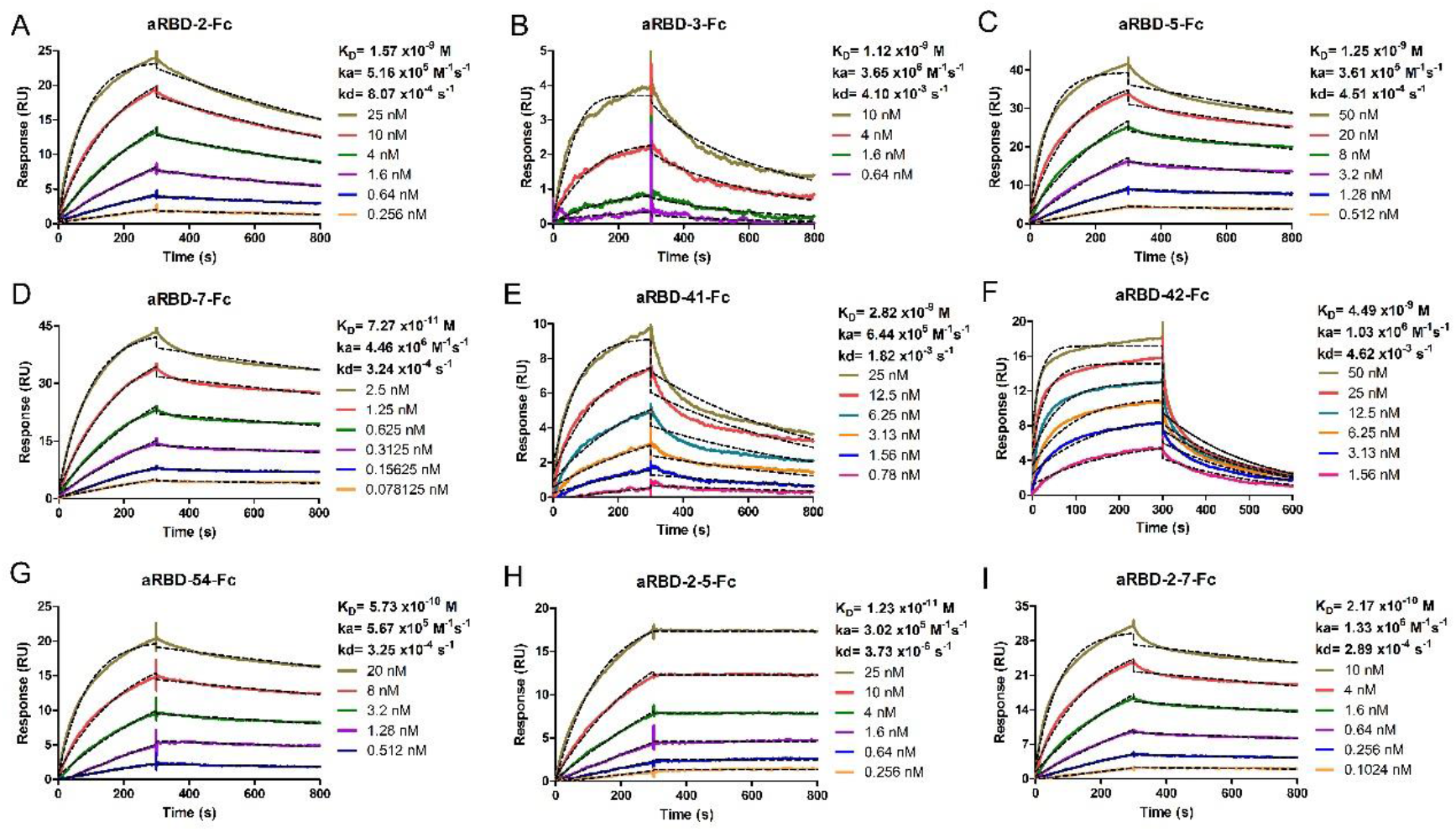
RBD-binding activity characterization of isolated Nb-Fc fusions using SPR. Binding kinetics of aRBD-2-Fc (A), aRBD-3-Fc (B), aRBD-5-Fc (C), aRBD-7-Fc (D), aRBD-41-Fc (E), aRBD-42-Fc(F), aRBD-54-Fc (G), aRBD-2-5-Fc (H) and aRBD-2-7-Fc (I) was measured by SPR. The SARS-CoV-2 RBD was immobilized onto a CM5 sensor chip, Nb-Fc fusions with serially 1:1 dilutions were injected and monitored by Biacore T200 system. Binding curves are colored lines, and fit of the data to a 1:1 binding model are black dotted lines.

**Fig. S7.**
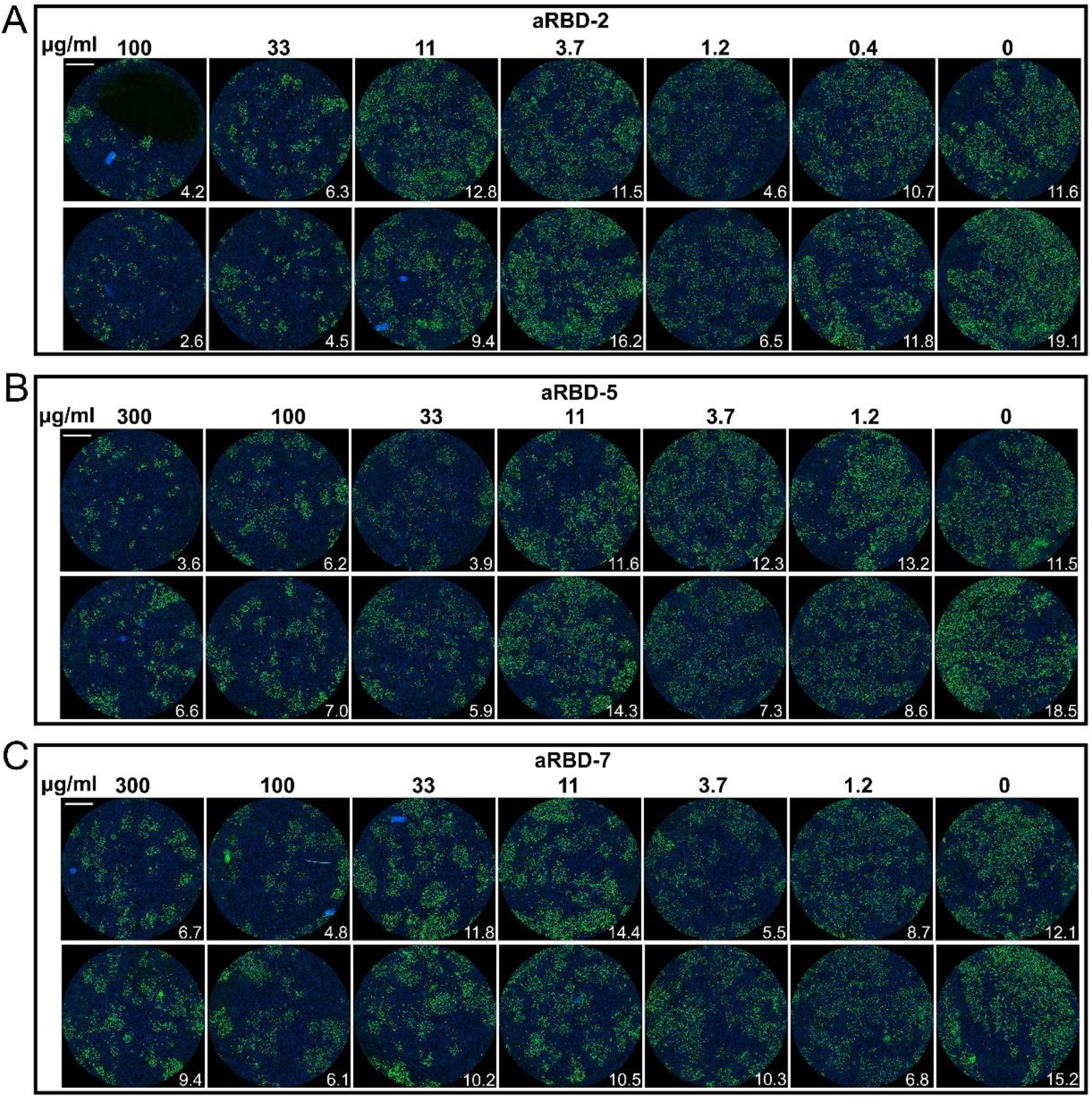
*In vitro* SARS-CoV-2 neutralization of three representative Nb monomers. The serially diluted aRBD-2 (A), aRBD-5 (B) and aRBD-7 (B) was incubated with ∼200 PFU of SARS-CoV-2. The mixture were then added to Vero E6 cells in 96-well plates. After 2 days of infection, the cells were stained with a monoclonal antibody against SARS-CoV-2 NP and an Alexa Fluor 488-conjugated goat anti-mouse secondary antibody. The nucleus was stained blue with Hoechst 33342. The experiment was performed in duplicate. Numbers in the low right corner of each grid were the percentage of cells infected by the virus.

**Fig. S8.**
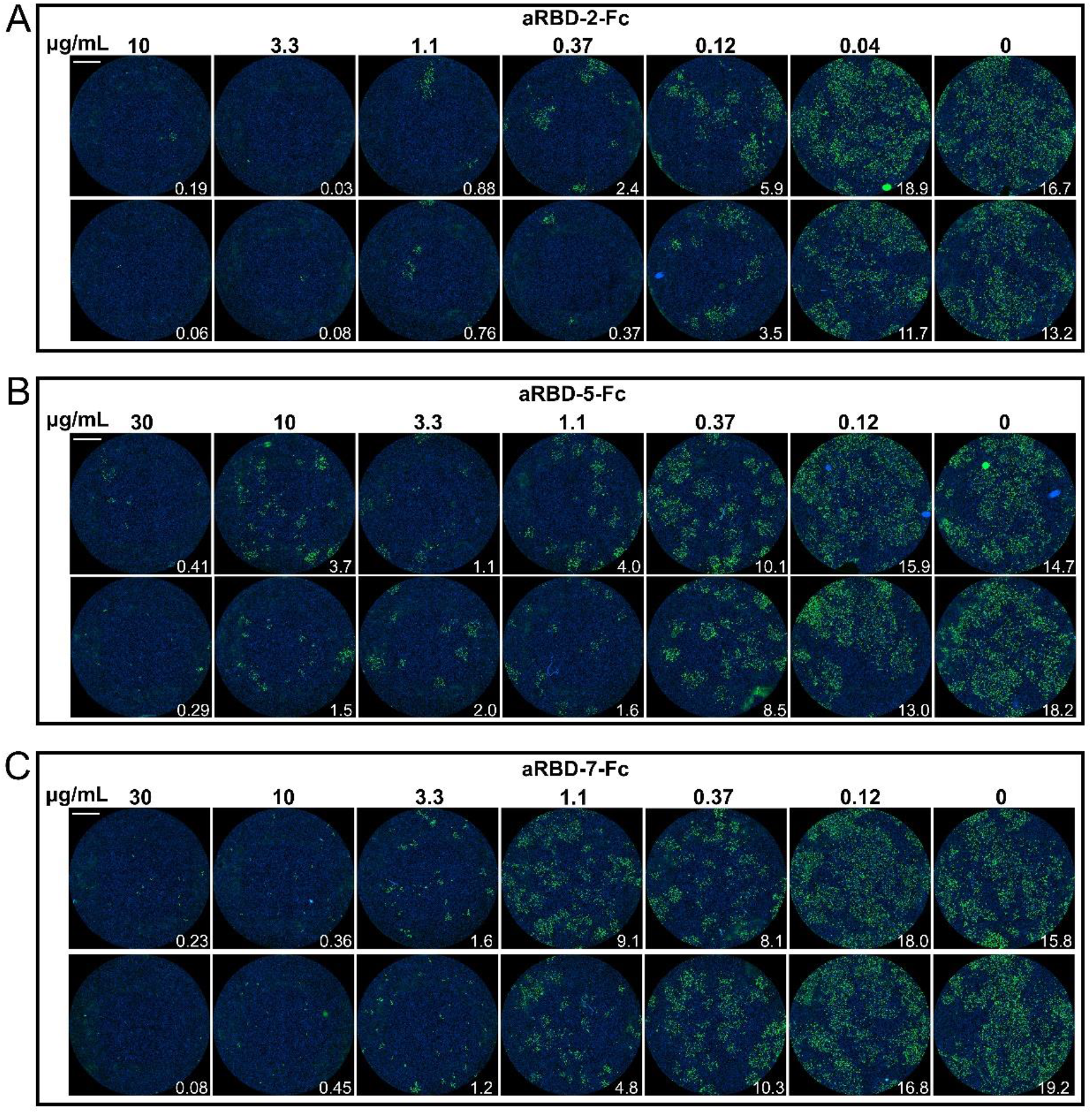
*In vitro* SARS-CoV-2 neutralization of Nb-Fc fusions. The serially diluted aRBD-2-Fc (A), aRBD-5-Fc (B) and aRBD-7-Fc (C) was incubated with ∼200 PFU of SARS-CoV-2. The mixture were then added to Vero E6 cells in 96-well plates. After 2 days of infection, the cells were stained with a monoclonal antibody against SARS-CoV-2 NP and an Alexa Fluor 488-conjugated goat anti-mouse secondary antibody. The nucleus was stained blue with Hoechst 33342. The experiment was performed in duplicate. Numbers in the low right corner of each grid were the percentage of cells infected by the virus.

## Notes

### Competing Interest Statement

The authors have declared no competing interest.

